# Method for identification of heme-binding proteins and quantification of their interactions

**DOI:** 10.1101/2019.12.17.879395

**Authors:** Nina Bozinovic, Rémi Noé, Alexia Kanyavuz, Maxime Lecerf, Jordan D. Dimitrov

## Abstract

The standard assay for characterization of interaction of heme with proteins is absorbance spectroscopy. However, this approach demands relatively large quantities of proteins and it is difficult to perform in high-throughput manner. Here, we describe an immunosorbent assay based on the covalent *in situ* conjugation of heme to a pre-coated carrier. Advantage of this assay is that it allows both identification of heme-binding proteins and quantification of their binding avidity, using only minimal amounts of protein (1-10 μg). Importantly, the same approach can be used for covalent linkage of other natural or synthetic compounds and analyzing their interactions with proteins.

## 1. Introduction

Heme is an indispensable constituent of hemoproteins (hemoglobin, myoglobin, cytochromes, peroxidases). Heme can also regulate functions of many cellular and plasma proteins by transient interactions [1, 2]. For example, it has been demonstrated that the normal immune repertoire contains antibodies that bind to heme and consequently acquire antigen binding polyreactivity [3, 4]. There is evidence suggesting that heme binds to the variable region of antibodies and serves as an interfacial cofactor for antigen recognition [4, 5]. In order to identify proteins capable of transient interaction with heme, UV-vis absorbance spectroscopy can be applied. This technique allows measurement of interaction between proteins and heme. However, absorbance spectroscopy is not appropriate for screening of large libraries of proteins; it is time-consuming and requires relatively large quantities of protein.

ELISA (enzyme-linked immunosorbent assay) is cost-efficient, high-throughput and technically simple to perform. It is widely used to determine binding of immunoglobulins to protein antigens. Proteins are easily coated on the polystyrene surface. However, being a small molecule (652 Da), heme does not possess sufficient molecular size for appropriate coating on the polystyrene surface. For their efficient immobilization, low molecular weight compounds (haptens) are usually conjugated to carrier proteins. Nonetheless, the preparation and purification of protein conjugates especially with hydrophobic haptens, such as heme, is a complex procedure and may not be routinely implemented in all laboratories.

Here, we describe an alternative experimental technique based on ELISA, where heme is covalently linked *in situ* to a carrier peptide pre-coated on the surface. This technique is fast, cost efficient and can be applied in high-throughput manner. Moreover, it allows the quantification and comparison of the binding avidity of heme-binding proteins. The assay can also be applied for the evaluation of the interaction of proteins with other synthetic or natural heterocyclic compounds.

## 2. Materials and methods

### 2.1. Materials

Hemin, Fe(III)-protoporphyrin IX chloride, was obtained from Frontier Scientific, Inc. (Logan, UT). 1-ethyl-3-(3-dimethylaminopropyl) carbodiimide hydrochloride (EDC) and *N*-hydroxysulfosuccinimide (Sulfo-NHS) were obtained from Thermo Fisher Scientific (Waltham, MA). DMSO was obtained from Sigma-Aldrich (St. Louis, MO). All chemicals were with the highest available purity. Human recombinant IgG1 antibodies cloned from B cells isolated from synovial tissue of rheumatoid arthritis patients (Ab15, Ab16, Ab17, Ab18, Ab19, Ab20, Ab21, Ab49, Ab56, Ab65, Ab72 and Ab137) were kindly provided by Prof. Claudia Berek (DRFZ, Berlin). N07G05 is a monoclonal IgG1 cloned from human naïve B cell. Heme-binding antibody Ab21 was thoroughly dialyzed against PBS containing 10% sucrose and stored before use at –20 °C at concentration of 12 mg/mL. Human pooled immunoglobulin G (IVIg, Endobulin, Baxter USA) was dialyzed against PBS and stored before use at –20 °C at concentration of 80 mg/mL. Hemin was dissolved in DMSO to final concentration of 10 mM. Human C1q was obtained from Calbiochem (Burlington, MA). Human hemoglobin was obtained from Sigma-Aldrich (St. Louis, MO). Preparation of apo-hemoglobin (apo-Hb) was done by procedure described in [6].

### 2.2. Immunosorbent assay for assessment of antibody binding to heme

#### 2.2.1. *In situ* conjugation of heme

Ninety-six-well polystyrene plates (Nunc MaxiSorp) were first coated with 100 μl /well of 0.5 % gelatin in PBS. For facilitating the solubilization of gelatin the solution was heated in water bath to 40-45 °C. After incubation for 1 h at room temperature, plates were washed (3 ×) with water. Solutions of 1 mM heme in solvent mixture DMSO/H_2_O = 1/1 was added to the microtitration plate at 50 μl /well. Next either: a) deionized water; b) 60 mM EDC in water, or c) 60 mM EDC and 60 mM Sulfo-NHS in water, were added to the heme-containing wells in the volume ratio 2:1 (i.e. 25 μl /well). The solutions were well homogenized. Plates were incubated at room temperature in dark with a gentle shaking for 2 h. After, the wells were washed (3 ×) with water, then incubated for 5 min with 1M water solution of ethanolamine at pH 8, and again washed (3 ×) with water.

#### 2.2.2. Assessment of binding of proteins to heme-conjugated surface

The residual binding sites on plates were blocked by incubation with PBS containing 0.25% Tween 20. For assessment of the antibody binding, Ab21 and Ab72 were diluted to 100 μg/mL (670 nM) with PBS-T (0.05% Tween 20) and were further serially diluted with PBS-T in the range of 100 − 0.195 μg/mL (670 – 1.3 nM, dilution factor of 2). Antibodies were incubated for 1 h at room temperature with plate coated with heme-modified gelatin. After incubation with the antibody, the plate was washed extensively (5×) with PBS-T and incubated with a HRP-conjugated mouse anti-human IgG (9040-05, clone JDC-10, Southern Biotech, Birmingham, AL) for 1 h at room temperature. Binding of Ab21 was revealed by measuring the absorbance at 492 nm after the addition of peroxidase substrate, *o*-phenylenediamine dihydrochloride (Sigma-Adrich) and stopping the reaction by the addition of 2 M HCl. Measurement of the absorbance was performed with a microplate reader (Infinite 200 Pro, Tecan, Männedorf, Switzerland).

Following sections describe variations of the procedure. Initial and final steps of the experiments are identical to those described above. For the covalent binding of heme to gelatin, activation of 1 mM heme was done with 60 mM EDC, unless stated otherwise.

#### 2.2.3. Analyses of the binding of pooled human IgG

For binding assessment, pooled therapeutic human IgG (IVIg) preparation, was first diluted to 1000 μg/mL (6.7 μM) in PBS-T (0.05% Tween 20) and was further serially diluted in the range of 1000 − 3.90 μg/mL (6.7 – 0.026 μM, dilution factor of 2). Antibodies were incubated for 1 h at room temperature with plate coated with heme-modified gelatin.

#### 2.2.4. Comparison of antibodies binding to heme

For binding assessment, a set of monoclonal antibodies were diluted to 10 μg/mL (67 nM) with PBS-T (0.05% Tween 20) and incubated (50 μl /well) for 1 h at room temperature with microtiter plate coated with heme-modified gelatin.

#### 2.2.5 Binding of C1q to heme

For binding assessment, C1q was diluted to 50 μg/mL (120 nM) with PBS-T (0.05% Tween 20) and was further serially diluted with PBS-T in the range, of 50 − 0.195 μg/mL (120 - 0.48 nM, dilution factor of 2). C1q was incubated for 1 h at room temperature with the plate coated with heme-modified gelatin. For detection of the binding of human C1q to conjugated heme, we used a sheep anti-human C1q conjugated with HRP (ab46191, Abcam, Cambridge, UK) diluted 1000 × in PBS-T. The plates were incubated with the antibody for 1 h at room temperature.

#### 2.2.6. Binding affinity of apo-hemoglobin

For binding assessment, human apo-hemoglobin was diluted to 100 μg/mL (1.5 μM) with PBS-T (0.05% Tween 20) and was further serially diluted with PBS-T in the range of 100 − 0.39 μg/mL (1.5 – 0.006 μM, dilution factor of 2). Apo-hemoglobin was incubated for 1 h at room temperature with the plate coated with heme-modified gelatin. After washing (5×) with PBS-T, the plates were incubated for 1 h at room temperature with goat polyclonal IgG to human hemoglobin (LS-B13233, LSBio, Seattle WA), diluted 1000 × in PBS-T. The immunoreactivity was detected by incubation with Rabbit F(ab’)2 Anti-Goat IgG(H+L)-HRP (6020-05, Southern Biotech) diluted 3000 × in PBS-T.

#### 2.2.7. Data analyses

The binding isotherms were obtained by non-linear regression analyses using GraphPad Prisms v.6 software (GraphPad Software, San Diego, CA). For fitting of the experimental data we applied the binding model that uses following equation: *Y=Bmax*X/(K*_*D*_*+X)+NS*X + Background*, where Bmax is the maximum specific binding; K_D_ is equilibrium binding constant that indicates the apparent affinity; NS is the slope of nonspecific binding; X is concentration of antibody or heme-binding protein, and Y is binding intensity. In case when we subtracted the background reactivity of antibodies towards unconjugated gelatin, we applied the following equation: *Y = Bmax*X/(Kd + X)* for fitting the data.

### 2.3. Absorbance spectroscopy

All absorbance spectroscopy measurements were performed with Cary-300 spectrophotometer (Agilent Technologies, Santa-Clara, CA). Proteins (human IgG1 antibodies – Ab21 and N07G05) were diluted to final concentration of 1.5 μM in PBS in volume of 1 ml. The dilution was done directly into quartz optical cells (Hellma, Jena, Germany) with 10 mm optical path. The antibodies were titrated with increasing concentrations of hemin (stock solutions in DMSO) resulting in final concentrations of 0.75, 1.5, 3, 6, 12 and 24 μM. Hemin aliquots were also added to an optical cell containing PBS only. Immediately after addition of hemin and intensive homogenization, the UV-vis spectra were recorded in the wavelength range 300-700 nm. All measurements were performed at room temperatures.

## Results and discussion

In order to identify and quantify interactions of proteins with heme, we developed an assay where heme is covalently bound to a carrier peptide coated on the plate surface. Here, we describe a procedure that was refined following testing different experimental conditions. This procedure resulted in the most efficient and reproducible immobilization of heme and detection of heme-binding proteins and therefore we recommend these conditions for analyses of protein-heme interactions.

Gelatin was chosen as carrier because it consists of smaller peptides, obtained by hydrolysis of 300 kDa protein collagen, hence reducing the possibility for non-specific interactions. Moreover, as gelatin represents protein hydrolysate, it does not have any defined tertiary structure that might be compromised by exposure to harsh conditions necessary for the conjugation reaction. Carboxyl groups of heme require chemical activation in order to be coupled with primary amino groups of lysine residues of peptides. Typical carboxyl activating agent EDC activates carboxylic acids by creating a more reactive ester leaving group. Less labile, but still a good leaving group can be formed when EDC is used in the combination with Sulfo-NHS. For the initial optimization of the assay, a human recombinant IgG1 Ab21 was chosen as a well characterized heme-binding antibody [5].

### 1) Preparation of binding surface

Polystyrene 96-well microtitration plates were coated for 1 hour with 0.5 % solution of gelatin in PBS. In order to achieve in situ heme conjugation to the carrier, after washing with deionized water, 1 mM solution of hemin in 50 % DMSO was added, followed by the addition of excess of the activating reagents – final concentration of 60 mM EDC or a mixture of 60 mM EDC and 60 mM Sulfo-NHS and incubation for 2 hours at room temperature. After, the wells were briefly incubated with 1M solution of ethanolamine at pH 8, in order to saturate possibly activated carboxyl groups on gelatin and to dissolve and wash away any non-reacted hemin. After the incubation with ethanolamine, the binding surface was blocked by exposure to 0.25 % Tween 20 in PBS.

To determine the speciation of heme while bound to gelatin we performed absorbance spectroscopy analyses. These analyses reveal that interaction of hemin with gelatin in the absence or presence of EDC did not result in major change in heme species observed in PBS (Supplemental Figure 1). This result indicate that hemin shows minimal non-specific interactions with the carrier and that the most probable species of heme displayed on the gelatin surface are heme dimers.

### 2) Assessment of protein-heme interactions

For binding assessment, increasing concentrations of Ab21 were incubated on the hemin-conjugated surface. Covalent coupling of hemin to gelatin was successful, which was indicated by the high intensity of Ab21 binding to gelatin exposed to hemin as compared to native gelatin (Figure 1A). Activation with EDC alone provided better results than the activation with the mixture of EDC and Sulfo-NHS (data not shown). It is possible that the less reactive intermediate formed with Sulfo-NHS requires longer reaction time or an increased temperature to react with primary amino groups from gelatin. The non-linear regression fit of the binding data allowed calculation of the apparent binding affinity (avidity) of the antibody to hemin. Thus, Ab21 recognized immobilized hemin with K_D_ of 56 ± 17 nM (based on estimation of K_D_ of Ab21 from the three independent experiments presented on Figure 1 panels A and B), suggesting that this antibody has relatively high binding avidity for heme.

**Figure 1.**
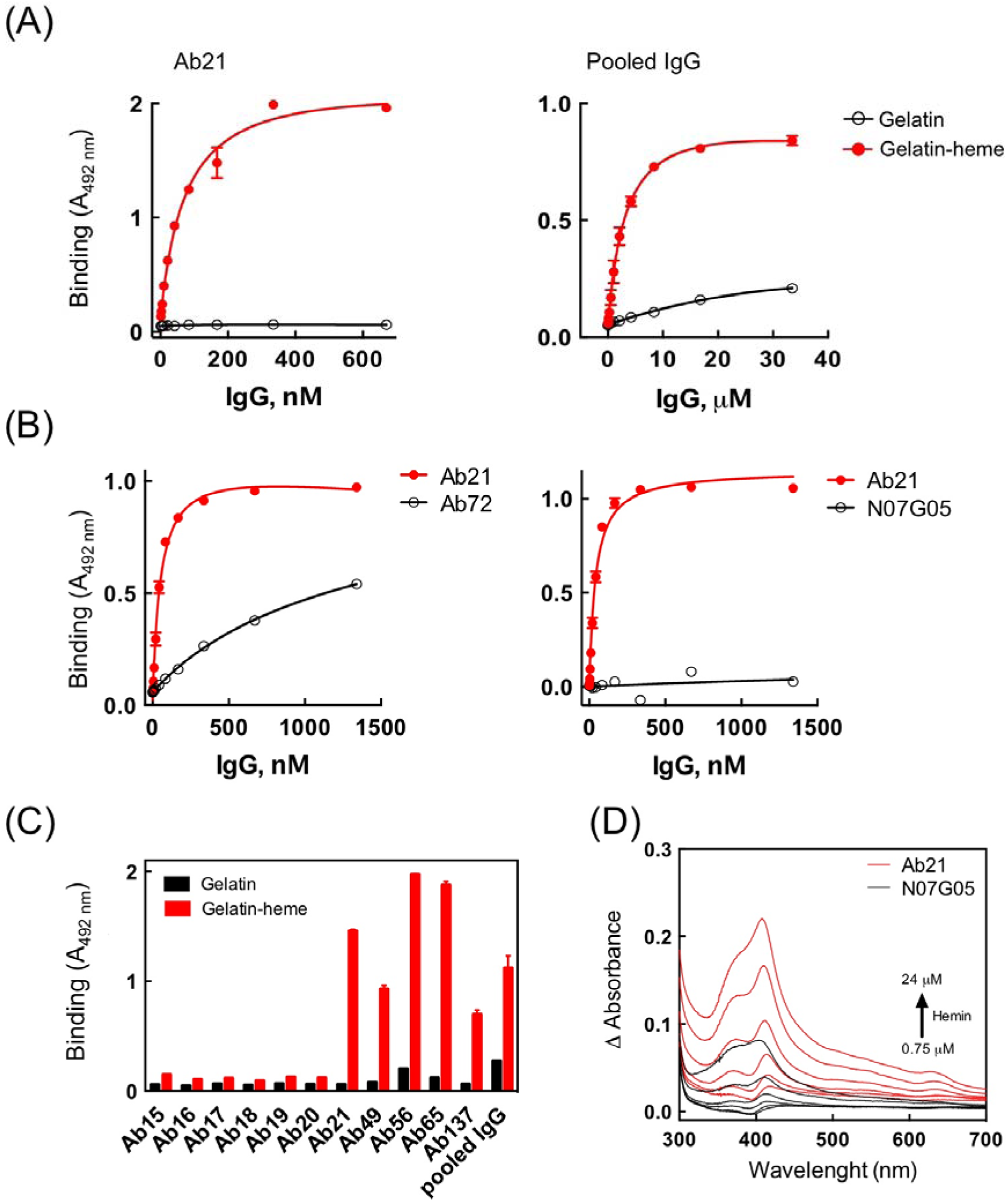
Analyses of interaction of antibodies with immobilized hemin. (A) Evaluation of binding of human heme-binding IgG1 Ab21 (left panel) or pooled human IgG1 to *in situ* peptide-conjugated hemin. The binding of antibodies to carrier (gelatin) only was also determined. The binding of each antibody to non-conjugated carrier was also assessed. (B) Comparison of the hemin-binding capacity of increasing concentrations of human recombinant IgG1, Ab21, Ab72 and N07G05. (C) Comparison of hemin-binding potential of a panel of human recombinant IgG1 antibodies and human pooled IgG. The monoclonal antibodies were tested at 10 μg/mL, pooled IgG was tested at 100 μg/mL. (D) Differential spectra in the wavelength range 300-700 nm, depicting the interaction of increasing concentration of hemin with antibodies Ab21 and N07G05. The mean binding intensity ±SD, obtained from three technical replicates are presented. Data presented on panels A and B are representative examples from one of two independent experiments.

Since a significant fraction of immunoglobulins in a human immune repertoire has been shown to interact with heme and acquire antigen binding polyreactivity [7, 8], it was expected that pooled IgG preparation obtained from plasma of large number of healthy donors would be able to bind to immobilized heme. To test this assumption, a commercially available pooled IgG preparation was essayed for hemin binding by the assay described here. The pooled human IgG exhibited a considerable heme-binding potential (Fig. 1A), confirming that normal immunoglobulin repertoire contain a fraction of heme-binding antibodies.

To further confirm that the assay can discriminate between heme-binding and heme-nonbinding monoclonal antibodies, binding of Ab21 to heme-modified gelatin was compared to the binding of other human monoclonal IgG1, Ab72 and N07G05. As can be observed in Figure 1B, the binding of Ab72 to heme was significantly lower than the binding of Ab21 in the same concentration range. N07G05 showed no specific binding to hemin.

The screening was further expanded to a set of human recombinant IgG1 antibodies that were pre-selected to have or not have potential to interact with heme. All antibodies were screened at a single concentration, 10 μg/mL (0.5 μg/well). The obtained results clearly demonstrated that the assay can well discriminate between the antibodies that were able to bind heme and those that did not (Figure 1C).

To confirm further the ability of the newly developed assay to discriminate between heme binding and non-binding proteins we applied absorbance spectroscopy (Figure 1D). These analyses revealed a clear difference in the differential absorbance spectra of the antibody identified as heme binding (Ab21) compared to the one that did not bind specifically to hemin (N07G05) (Figure 1D).

### 3) Validation of assay with apo-hemoglobin and C1q

To further validate the utility of the method for assessing the typical heme-binding proteins, we used apo-hemoglobin i.e. hemoglobin that lacks its prosthetic group. The apo-hemoglobin showed concentration dependent binding to the immobilized hemin (Figure 2A). The estimated apparent K_D_ value was 182 ± 55 nM. Further, the test was also applied for study of the interaction of non-conventional heme-binding protein – complement C1q [9]. This protein also demonstrated clear binding to surface immobilized heme with apparent K_D_ value of 21.4 ± 5.6 nM (Figure 2B). By using solution-based methods it was estimated that hemin binds to apo-Hb with binding constant (Kb) or 2.72 × 10^6^ equal to K_D_ value of 367 nM [10]. The apparent K_D_ value of the binding of hemin to C1q was lower (i.e. 1-2 μM) as estimated by solution based techniques [9]. The fact that in the present study C1q binds hemin with higher apparent binding affinity than apo-Hb can be explained by avidity effects. Of note C1q has six identical globular heads that are responsible for target recognition. The cooperative effect of multivalent interaction of C1q to surface immobilized heme can explain the estimated high apparent affinity.

**Figure 2.**
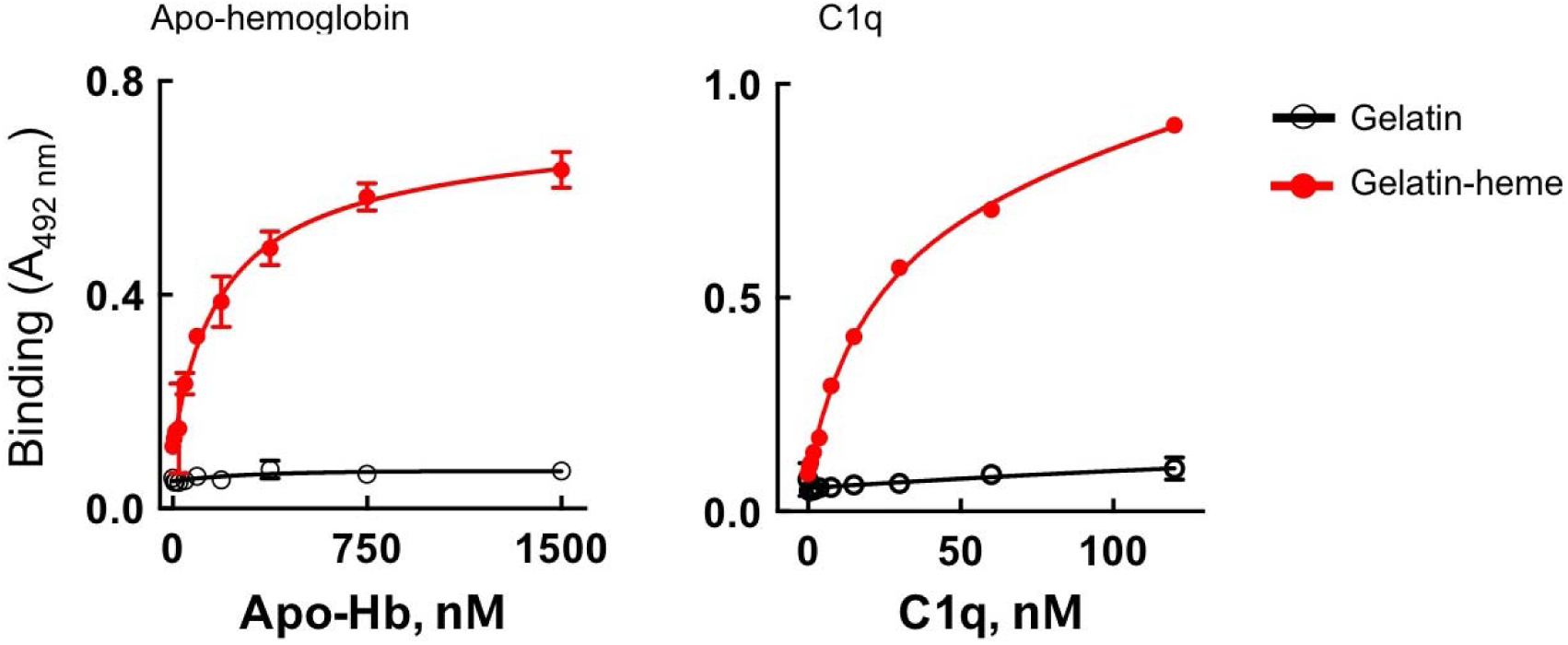
Analyses of interaction of apo-hemoglobin and C1q with immobilized heme. Binding of increasing concentrations of human apo-hemoglobin (A) and of human C1q (B) to hemin-conjugated to gelatin. The binding of the proteins to carrier (gelatin) only was also determined. The mean binding intensity ±SD, obtained from three technical replicates are presented. Data presented are representative examples from one of two independent experiments.

The present technique uses oxidized form of heme (hemin) as a ligand. Of note, some proteins interact with considerably higher affinity to reduced heme. We expect that the assay can be easily optimized and performed in the presence of agents reducing hemin (such as sodium dithionite), thus broadening its scope.

## 3. Concluding remarks

Here, we present a procedure that can be used for identification of heme-binding proteins. The proposed technique is simple, requires low quantities of protein and can be performed in high-throughput manner. In addition, it allows estimation of the binding avidity of the heme-binding proteins. The technique can be coupled with mass spectrometry for identification of high affinity heme-binding proteins in complex mixtures, as for example lysates from eukaryotic cells or bacteria.

## Acknowledgments

This work was supported by Institut National de la Santé et de la Recherche Médicale (INSERM, France), Centre National de la Recherche Scientifique (CNRS, France).

## Supplementary information

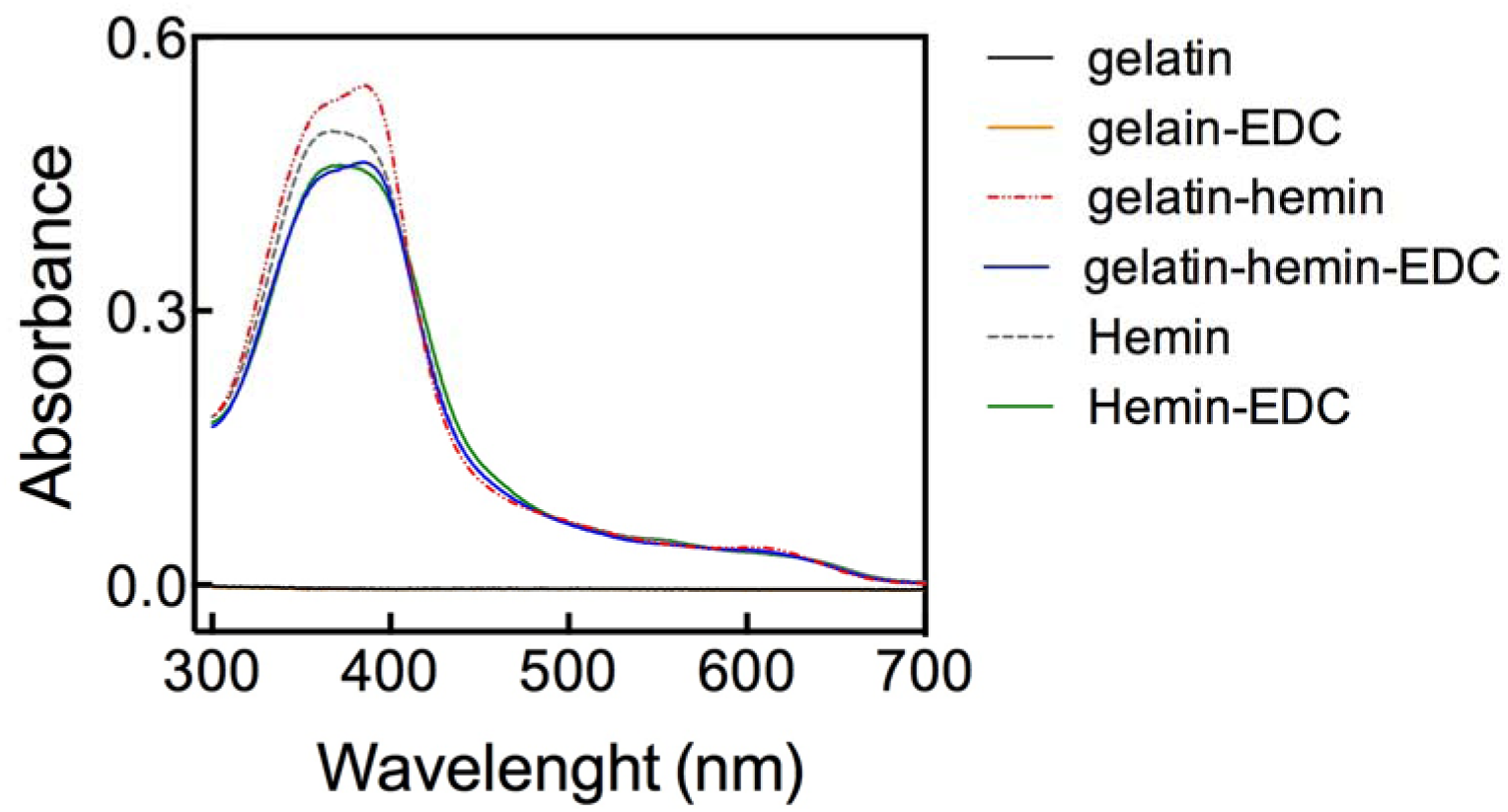

Absorbance spectroscopy. The gelatin at 0.05%, hemin at 100 μM and EDC at 2mM all diluted in PBS were incubated alone or in indicated combinations for 1h at room temperature. After the samples were diluted 10 × in PBS and UV-vis spectra were recorded in the range 300-700 nm. Of note, the lines depicting the absorbance of gelatin and gelatin-EDC are overlapping.

## Notes

### Competing Interest Statement

The authors have declared no competing interest.

### Summary of Updates

Material and methods updated. New results incorporated and discussed. Two new panels to Figure 1 added. New supplementary figure added.

